# Sequencing depth and genotype quality: Accuracy and breeding operation considerations for genomic selection applications in autopolyploid crops

**DOI:** 10.1101/2020.02.23.961383

**Authors:** Dorcus C Gemenet, Hannele Lindqvist-Kreuze, Bode A Olukolu, Bert De Boeck, Guilherme da Silva Pereira, Marcelo Mollinari, Zhao-Bang Zeng, G Craig Yencho, Hugo Campos

## Abstract

The autopolyploid nature of potato and sweetpotato ensures a wide range of meiotic configurations and linkage phases leading to complex gene action and pose problems in genotype data quality and genomic selection analyses. We used a 315-progeny biparental population of hexaploid sweetpotato and a diversity panel of 380 tetraploid potato, genotyped using different platforms to answer the following questions: i) do polyploid crop breeders need to invest more for additional sequencing depth? ii) how many markers are required to make selection decisions? iii) does considering non-additive genetic effects improve predictive ability (PA)? iv) does considering dosage or quantitative trait loci (QTL) offer significant improvement to PA? Our results show that only a small number of highly informative single nucleotide polymorphisms (SNPs; ≤ 1000) are adequate for prediction, hence it is possible to get this number at the current sequencing depth from most service providers. We also show that considering dosage information and additive-effects only models had the best PA for most traits, while the comparative advantage of considering non-additive genetic effects and including known QTL in the predictive model depended on trait architecture. We conclude that genomic selection can help accelerate the rate of genetic gains in potato and sweetpotato. However, application of genomic selection should be considered as part of optimizing the entire breeding program. Additionally, since the predictions in the current study are based on single populations, further studies on the effects of haplotype structure and inheritance on PA should be studied in actual multi-generation breeding populations.

**Key message:** Polypoid crop breeders do not need more investment for sequencing depth, dosage information and fewer highly informative SNPs recommended, non-additive models and QTL advantages on prediction dependent on trait architecture.

## Introduction

Phenotyping under recurrent selection has been an important approach for variety development in plant breeding, with substantial success to date. However, this process may take a long time for most crops, particularly for clonally propagated crops (**Slater et al. 2016**). For example, in potato, it typically takes an entire year to develop enough tubers from botanical seed obtained from crossing nurseries, for experimental trial purposes. This is followed by at least two years of field evaluation for qualitative traits, with evaluation for most quantitative traits in replicated multi-environment trials beginning in around year four (**Endelman et al. 2018**). The same can be said for sweetpotato, although cycle times in sweetpotato are shorter by about a year due to the fact that the crop can be vegetatively propagated via stem cuttings (**Wolfgang et al. 2009**). This represents a stark contrast with what can be achieved in cereal and legume crops, where up to 6 generations can be raised within a calendar year (**Watson et al. 2018**), or in private corn breeding programs based in the United States and Europe which can raise multiple generations per year through the coordinated use of winter nurseries located in both hemispheres such as United States, Puerto Rico, Hawaii and Chile. This therefore implies that the estimation of parental value based on genetic designs and phenotypic evaluation in potato and sweetpotato increases the selection cycle time, thereby reducing the rate of genetic gains and the speed of delivery of superior, novel genetics to farmers.

The use of genetic markers for selection offers potential to reduce the breeding cycle time as selection can be done at an earlier stage. Previously proposed methods have involved identifying quantitative trait loci (QTL) via QTL mapping and genome-wide association studies (GWAS), but they have had little practical application in the actual development of new cultivars through plant breeding to date, especially for complex quantitative traits, since identifying the causal genes underlying QTL needed to make their application practical is costly (**Xu and Crouch 2008**). Genomic selection (GS) offers the ability to select parents within a shorter interval and increase selection intensity by predicting untested genotypes earlier and enhancing larger starting genetic variation. This approach uses genome-wide marker data to predict the performance of untested genotypes and estimate their breeding values (genomic estimated breeding values; GEBVs), based on a genotyped and phenotyped training population **(Meuwissen et al. 2001)**. Genomic selection is emerging as the approach of choice to circumvent the limitations associated with use of QTL for marker-assisted selection and to improve the efficiency of phenotypic selection (**Bernal-Vasquez et al. 2014**). Good genetic progress can be made using GS, as long as factors that affect its predictive ability (PA), i.e. the correlation between phenotypic best linear unbiased estimators (BLUPs) and GEBVs, are well understood. These include trait architecture, the size of the training population, the relationship between the training and validation populations, heritability of the trait, the quality of phenotypic efforts, the level of linkage disequilibrium (LD), marker density, environmental variances and covariance among traits (**Covarrubias-Pazaran et al. 2018**).

The application of GS is taking shape in plant breeding with more and more crops exploring its utility (**Spindel et al. 2016; Wang et al. 2018; Endelman et al. 2018; Covarrubias-Pazaran et al. 2018; Faville et al. 2018; Nyine et al. 2018; Bhandari et al. 2019).** For crops like rice and wheat that are normally self-pollinated and have a high incidence of high-effect QTL (**Spindel et al. 2016**), faster success is expected from applying GS as prediction accuracy depends primarily on the factors listed above. However, breeders of auto-polyploid, clonally propagated crops like potato and sweetpotato, which are normally heterogenous and heterozygous, have to ask themselves additional questions and identify trade-off points that enhance the success of GS-assisted breeding (**Slater et al. 2016; Endelman et al. 2018**). Potato and sweetpotato present a wide range of meiotic configurations and linkage phases (**Mollinari et al. 2020**). In addition to causing complex gene action effects, allelic and configuration diversity have consequences on genotyping and genotype data quality, which consequently affects downstream analysis for quantitative-genetic parameters required to make high quality breeding decisions. Genotyping-by-sequencing (GBS) has currently become a genotyping method of choice in plant breeding (**Poland and Rife 2012**) but it is also prone to genotyping errors and a high level of missingness at low depth of sequencing, while high sequencing depth has additional cost implications. Data from polyploid crops is more prone to low quality genotype calls at low sequencing depth when compared to diploid crops, because of uncertain allele dosages and possibility of non-random inheritance of alleles such as in preferential pairing or double reduction (**Blischak et al. 2016, 2018**).

Public sector breeding programs like those conducted in centers which are part of the Consultative Group on International Agricultural Research (CGIAR), and in the individual National Agricultural Research Systems (NARS) existing in many countries, are currently undergoing breeding program optimization efforts in order to keep up with the challenges of climate change and population increase (**Cobb et al. 2019**). Application of GS is one such tool for breeding program optimization. In order to develop GS tools to make more effective breeding efforts in auto-polyploid crops such as potato and sweetpotato, we have taken a practical perspective within a plant breeding setting to address several pertinent questions related to application of GS in auto-polyploids. We used real data sets from a 380 training-panel made up of advanced tetraploid potato clones and a 315-full-sib family of hexaploid sweetpotato, both developed by the International Potato Center (CIP) and genotyped using different platforms, to address the following questions: i) do polyploid crop breeders need to invest more resources for additional sequencing depth? ii) how many genetic markers are required to make selection decisions? iii) does the consideration of non-additive genetic effects add value to predictive ability (PA) to enhance genetic gains either for population improvement or product development in polyploid crops? iv) given the multiple alleles at loci with diverse meiotic configurations and linkage phases, does considering dosage, haplotypic or QTL effects offer significant improvement to PA to enhance genetic advances? We also discuss other factors that need to be considered while adopting GS as a decision support tool in an optimized breeding program.

## Materials and Methods

### Genetic Materials and Phenotyping

#### Sweetpotato bi-parental population

A wide genetic variability exists in sweetpotato in terms of yield, nutritional content and culinary aspects, abiotic stress tolerance, biotic stress tolerance, among other attributes (**Low et al. 2017**). Introgression of high β-carotene content into locally adapted varieties is a major breeding objective especially in sub-Saharan Africa where vitamin A deficiency is prevalent. A 315-progeny full-sib family was developed by crossing a US-bred high β-carotene variety, ‘Beauregard’, with an adapted, locally preferred, starchy, low β-carotene landrace variety, ‘Tanzania’, at CIP-Peru. These two parents differ in additional traits of interest and the population will henceforth be referred to as the BT population. The population was evaluated in six environments of Peru, and six environments of Uganda for various quality-related and yield-related traits as described by **Gemenet et al. (2020)** and **Pereira et al. (2019**), between 2016 and 2017. The full 315-progeny population was also genotyped together with the parents using an optimized protocol for hexaploid sweetpotato, ‘GBSpoly’ at North Carolina State University (NCSU). To support genotyping protocol optimization for the hexaploid, a diploid relative of sweetpotato, *Ipomoea trifida*, was used to develop a full-sib family of about 200. The family was developed from two *I. trifida* lines referred to as M9 and M19, hence the M9 x M19 population, and also genotyped at NCSU. Additionally, a sub-sample of 292-progeny and the two parents of the BT population were genotyped by DArTSeq^TM^ in Australia, under the collaboration between the Integrated Genotyping Service and Support (IGSS) platform at the Biosciences east and central Africa (BecA) hub in Nairobi, Kenya and DArT. The quality-related traits measured in the BT population include: dry matter (DM) content, measured as a percentage of the laboratory dried samples divided by the initial fresh weight of 100g; Starch and β-carotene (BC) content, estimated using near-infrared reflectance spectroscopy (NIRS), and flesh color (FC), measured using internal color scales developed by CIP and partners. All quality-related traits were measured in Peru, but only flesh color was measured in Uganda (FC_U). Data is further described in **Gemenet et al. (2020**). For yield-related traits, total number of storage roots (TNR), number of commercial storage roots (NOCR), weight of total storage roots (RYTHA), weight of commercial storage roots (CYTHA), and total weight of foliage (FYTHA), were measured in the six experiments of Peru. Data is further described in **Pereira et al. (2019)**. Trait abbreviations are further defined in Table 1.

**Table 1.** Trait abbreviations and their description as used in the current study

| Crop | Trait Abbreviation | Trait Description |
| --- | --- | --- |
| Sweetpotato | DM | Dry matter content |
|  | Starch | Starch content |
|  | BC | Beta-carotene |
|  | FC_P | Flesh color in Peru |
|  | FC-U | Flesh color in Uganda |
|  | NOCR | # commercial storage roots |
|  | TNR | # total storage roots |
|  | CYTHA | Commercial storage root weight |
|  | RYTHA | Total storage root weigh |
|  | FYTHA | Total foliage yield weight |
| Potato | LB2014_O | Late blight in 2014 in Oxapampa, Peru |
|  | LB2016_Y | Late blight 2016 in Yunnan, China |
|  | PVY_L | Potato virus Y in Lima, Peru |
|  | AYP_K | Average yield per plant in Kunming, China |
|  | WMT_K | Weight of marketable tubers in Kunming, China |
|  | TTW16_Ica | Total tuber weight in 2016 in Ica-Peru |
|  | TTW16_HLJ | Total tuber weight in 2016 in Heilongjinag, China |

The quality-related traits were analyzed by fitting the following linear mixed model in ASREML:

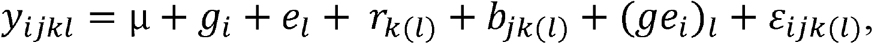

where *y*_*ijkl*_ = the vector of phenotypes of the individual *i* in block *j* within replicate *k* of environment *l*, *µ* = population mean, *g*_*i*_ the fixed treatment (genotype) effect, *e*_*l*_ = the random effect of environment *l*, *r*_*k(l)*_ = random effect of replicate *k* in environment *l*, *b*_*jk(l)*_ = random effect of block *j* within replicate *k* of environment *l*, (*ge*_*i*_)_*l*_ = random effect of individual i in environment l (L=5) environments, *ε*_*ijk(l)*_ = random error of the residuals, assuming 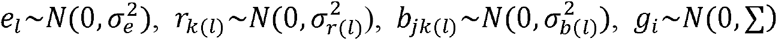 with ∑ = variance-covariance matrix across L environments, which varies according to the trait, 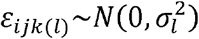 (**Gemenet et al. 2020**).

The yield-related traits were also analyzed with linear mixed models as described by **Pereira et al. (2019**) using GENSTAT 14 as:

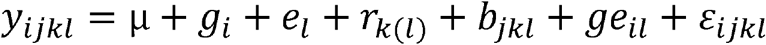

where *y*_*ijkl*_= the vector of phenotypes as above, μ=population mean, *g*_*i*_ the fixed treatment (genotype) effect, *e*_*l*_ = fixed effect of environment l, *r*_*k(l)*_ = fixed effect of replication k in environment l, *b*_*jkl*_ = random effect of block j within replication k in environment l; 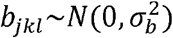, *ge*_*il*_ = the fixed interaction effect of individual i and environment l, and 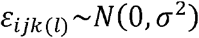 is the random residual error. The best linear unbiased estimators (BLUEs) as obtained by fitting the above models to the experimental data were then used to estimate GEBVs

#### Potato trait observation network population

A 380-genotype panel made up of advanced clones from the potato breeding program and representing all breeding populations at CIP was assembled for a trait observation network (TON) in Peru, China and Ethiopia. Henceforth, we shall refer to this population as the TON panel. The evaluation of the panel was carried out in diverse agro-ecological zones, and in subsets of genotypes subject to participating NARS’ partner capacity and/or ability to produce enough mini-tubers for experimentation. The experimental sites, experimental designs and the number of genotypes evaluated per experiment are summarized in Table 2. The TON panel was evaluated for maturity (bulking) by tuber characteristics at three harvest dates where average yield per plant (kg; AYP), weight of marketable tubers per plant (kg; WMT), were measured. Additionally, mature tuber weight was evaluated by measuring total tuber weight per plant (TTW; kg). In Peru, TTW was measured as the average total tuber weight across three drought-related treatments: terminal drought (irrigation stopped at flowering until harvest; TTW16_TD), recovery (partially irrigated after drought stress; TTW16_REC), and fully irrigated (normally irrigated throughout the growth period; TTW16_NI), while random drought was used in China, with no controlled treatments. Resistance to potato virus Y (PVY) was evaluated after infection with virulent vectors and susceptible spreader rows using standard protocols at CIP, while late blight resistance (LB) was evaluated by growing the population in endemic disease pressure and scored using standard protocols at CIP. Trait abbreviations are defined in Table 1. Additionally, the 380 genotypes were genotyped by GBS at Cornell University.

**Table 2.**
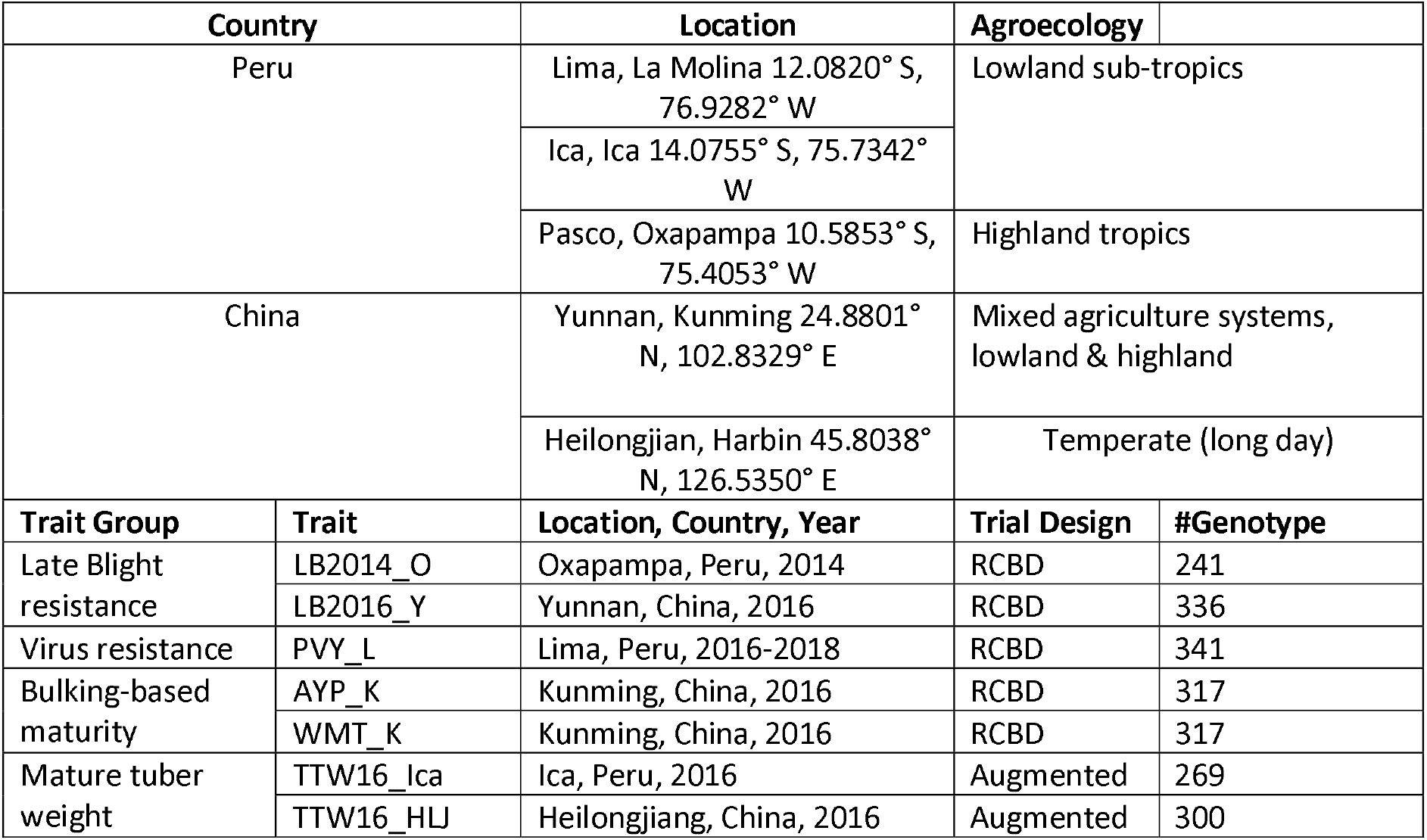
Locations, designs and traits measured in the trait observation network (TON) panel of potato

| Country |  | Location | Agroecology |  |
| --- | --- | --- | --- | --- |
| Peru |  | Lima, La Molina 12.0820° S,<br>76.9282° W | Lowland sub-tropics |  |
|  |  | Ica, Ica 14.0755° S, 75.7342°<br>W |  |  |
|  |  | Pasco, Oxapampa 10.5853° S,<br>75.4053° W | Highland tropics |  |
| China |  | Yunnan, Kunming 24.8801°<br>N, 102.8329° E | Mixed agriculture systems,<br>lowland & highland |  |
|  |  | Heilongjian, Harbin 45.8038°<br>N, 126.5350° E | Temperate (long day) |  |
| Trait Group | Trait | Location, Country, Year | Trial Design | #Genotype |
| Late Blight<br>resistance | LB2014_O | Oxapampa, Peru, 2014 | RCBD | 241 |
|  | LB2016_Y | Yunnan, China, 2016 | RCBD | 336 |
| Virus resistance | PVY_L | Lima, Peru, 2016-2018 | RCBD | 341 |
| Bulking-based<br>maturity | AYP_K | Kunming, China, 2016 | RCBD | 317 |
|  | WMT_K | Kunming, China, 2016 | RCBD | 317 |
| Mature tuber<br>weight | TTW16_Ica | Ica, Peru, 2016 | Augmented | 269 |
|  | TTW16_HLJ | Heilongjiang, China, 2016 | Augmented | 300 |

The experiments were analyzed as single trials, depending on the experimental design used as summarized in Table 2. A linear mixed model, taking into account the respective experimental design, was fitted to the phenotypic data. For those traits with different treatments like TTW in Peru, the joint adjusted means were additionally obtained across all treatments by fitting a linear mixed model. Genotype was considered as a fixed effect in these mixed models, so that BLUEs for the genotypic means were obtained for each trait and used to predict GEBVs.

### Genotyping and Variant Calling

#### DArTSeq^TM^ for Sweetpotato

DArTseq™ represents a combination of DArT complexity reduction methods and next generation sequencing platforms (**Kilian et al. 2012; Courtois et al. 2013; Raman et al. 2014; Cruz et al. 2013**). Therefore, DArTseq™ represents a new implementation of sequencing complexity reduced representations (**Altshuler et al. 2000**) and more recent applications of this concept on the next-generation sequencing platforms (**Baird et al. 2008; Elshire et al. 2011**). Similar to previous DArT methods based on array hybridizations, the technology is optimized for each organism and application by selecting the most appropriate complexity reduction method (both the size of the representation and the fraction of a genome selected for assays). Four methods of complexity reduction were tested in sweetpotato (data not presented) and the *PstI-MseI* method was selected. DNA samples were processed in digestion/ligation reactions principally as per **Kilian et al. (2012)** but replacing a single *PstI*-compatible adaptor with two different adaptors corresponding to two different Restriction Enzyme (RE) overhangs. The *PstI*-compatible adapter was designed to include Illumina flowcell attachment sequence, primer sequence and “staggered”, varying length barcode region, similar to the sequence reported by **Elshire et al. (2011**). This reverse adapter contained a flowcell attachment region and a *MseI*-compatible overhang sequence. Only “mixed fragments” (*PstI-MseI*) were effectively amplified in 30 rounds of PCR using the following reaction conditions: i) 94□ C for 1 min, ii) 30 cycles of: 94□ C for 20 sec, 58□ C for 30 sec, 72□ C for 45 sec, iii) 72□ C for 7 min. After PCR, equimolar amounts of amplification products from each sample of the 96-well microtiter plate were bulked and applied to c-Bot (Illumina) bridge PCR followed by sequencing on Illumina Hiseq2000. The sequencing (single read) was run for 77 cycles. Sequences generated from each lane were processed using proprietary DArT analytical pipelines. In the primary pipeline the FastQ files were first processed to filter away poor-quality sequences, applying more stringent selection criteria to the barcode region compared to the rest of the sequence. This was to ensure reliability in the assignments of the sequences to specific samples carried in the “barcode split” step. Approximately 2,000,000 sequences per barcode/sample were identified and used in marker calling. Finally, identical sequences were collapsed into “fastqcoll files”. The fastqcoll files were “groomed” using DArT PL’s proprietary algorithm which corrects low quality base from singleton tag into a correct base using collapsed tags with multiple members as a template. The “groomed” fastqcoll files were used in the secondary pipeline for DArT PL’s proprietary SNP and SilicoDArT (presence/absence of restriction fragments in representation) calling algorithms (DArTsoft14). For SNP calling, all tags from all libraries included in the DArTsoft14 analysis were clustered using DArT PL’s C++ algorithm at the threshold distance of 3, followed by parsing of the clusters into separate SNP loci using a range of technical parameters, especially the balance of read counts for the allelic pairs. Additional selection criteria were added to the algorithm based on analysis of approximately 1,000 controlled cross populations. Testing a range of tag counts parameters facilitated selection of true allelic variants from paralogous sequences. In addition, multiple samples were processed from DNA to allelic calls as technical replicates and scoring consistency was used as the main selection criteria for high quality/low error rate markers. Calling quality was assured by high average read depth per locus (>30X). The SNPs were coded as 0 = AA, 1 = BB, 2 = AB and “-“ = Missing. The sequences were not aligned to a reference genome because by the time of genotyping, the diploid references (**Wu et al. 2018**) had not been published.

#### GBSPoly© for Sweetpotato

GBSpoly is an optimized protocol for hexaploid sweetpotato developed at NCSU as part of a project focusing on developing genomic tools for sweetpotato improvement. The DNA was checked for quality on 1% agarose gel and quantified based on the PicoGreen florescence-based assay and the concentration was normalized to 50 ng/µl. Initially, several optimization efforts regarding restriction enzyme pairing was carried out (data not shown) and *CviAII-TseI* was selected to be the best combination for hexaploid sweetpotato. Therefore, 1µg of DNA was double-digested using five units of *CviAII* for three hours at 25°C followed by digestion with *TseI* for another three hours at 65°C. A new England Biolabs (NEB) CutSmart buffer was used to make up a total volume of 30 µl. Purification of the digested samples was done using AMPure XP magnetic beads from ThermoFisher^TM^ and quantified with PicoGreen assay. Barcodes were designed to account for substitution and indel errors and had an 8-bp buffer sequence to ensure that the barcode lay within high-quality base call regions of the sequence reads. Additional double digests on 64-plex pooled samples, purification, and size selection steps were carried out as described by **Wadl et al. (2018)** before performing 125 bp single-end sequencing on a total of 40 sequencing lanes (8 lanes for each of the 5 libraries) of the Illumina HiSeq 2500 platform. The resultant FastQ files were aligned to reference genomes of two wild relatives of sweetpotato, *Ipomoea trifida* and *Ipomoea triloba*, and variant calling done using the GBSapp pipeline as described by **Wadl et al. (2018)**. The SNPs were coded according to the dosage of the alternative allele as 0 = AAAAAA, 1 = AAAAAB, 2 = AAAABB, 3 = AAABBB, 4 = AABBBB, 5 = ABBBBB, 6 = BBBBBB. The variant calling process is summarized in **Online Resource 1**.

#### GBS-Cornell for Potato

The 380-genotype TON panel was genotyped by Cornell University using GBS in 2015. The DNA was digested with *EcoT221* restriction enzyme and 48-plex libraries were prepared for sequencing, using customized GBS protocols at Cornell. The resultant FastQ files were quality controlled and variant calling done using GATK HaplotypeCaller option (**Poplin et al. 2017**), disabling the duplicate read filter (this is recommended for GBS data) and using the joint genotyping -ERC GVCF mode. The reads were aligned to the potato genome reference sequenced from *S. tuberosum* group Phureja, line DM1-3 516 R44, a doubled monoploid (DM) via anther culture by the potato genome sequencing consortium (PGSC). Version PGSC_DM_v4.03 of the reference genome was used in alignment. The barcodes were removed using stacks and the ends were trimmed using trim-galore, followed by mapping to the reference using BWA. Resultant SAM files were processed using samtools and variants called using GATK Haplotype caller, targeting biallelic SNPs only. The SNPs were coded according to the dosage of the alternative allele as 0 = AAAA, 1 = AAAB, 2 = AABB, 3 = ABBB, and 4 = BBBB. The SNP filtration was done using bcftools allowing only for those SNPs with MAF of ≥ 3%, missingness of ≤ 2%, average genotype quality (GQ) ≥ 20 and average allele sequencing depth (DP) ≥ 16.

### Model comparison for predictive ability

We used the AGHmatrix package (**Amadeu et al. 2016**) to develop kinship G-matrices partitioning genetic variation based on several gene-action models. For the BT population DArTSeq markers (sweetpotato) where we did not have dosage information, we developed an additive G-matrix according to **VanRaden (2008)**, here-in referred to as Add_2x_DArTseq, and a non-additive effects G-matrix according **Vitezica and colleagues (2013)**, herein referred to as NonAdd_2x_DArTSeq. For the BT population GBSpoly (sweetpotato) and TON population GBS-Cornell (potato) data where we had dosage information, we employed three models to develop the G-matrices: (i) modeling only additive effects, according to **VanRaden (2008)** here-in referred to as Add_6x_GBSpoly for sweetpotato and Add_4x_GBSCornell for potato, (ii) modeling additive plus non-additive effects, according to **Slater et al. (2016)** here-in referred to as Add+Non_6x_GBSpoly for sweetpotato and Add+Non_4x_GBSCornell for potato, and (iii) a pseudo-diploidized effect model according to **Slater et al. (2016)**, here-in referred to as Pseudo_2x_GBSpoly for sweetpotato and Pseudo_2x_GBSCornell. The pseudo-diploidization collapses all dosage classes between the nulliplex and the hexaplex (in sweetpotato), and between the nulliplex and tetraplex (in potato) into one heterozygous class, under the assumption that all heterozygotes have an equal effect which falls in between both homozygotes. In the case of potato, the design matrix coding for the pseudo-diploid, additive autotetraploid and full autotetraploid was as described by **Slater et al. (2016)**, while that for sweetpotato is shown in Table 3. During kinship matrix development, additional filters were applied to the genotype data, to have MAF ≥ 30%, and missing data ≤ 10%. We used genomic best linear unbiased prediction (G-BLUP; **Clark and van der Werf 2013**) to compare the predictive ability of the five models for sweetpotato and three models for potato using the kinship matrices as variance-covariance matrices to fit the compressed linear mixed model (**Zhang et al. 2010**) and estimate genomic best linear unbiased predictors (G-BLUPs). The software GAPIT (**Lipka et al. 2012**) was used in the G-BLUP prediction fitting the following general model:

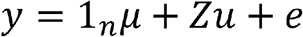

where y = vector of phenotypic data, 1_n_ is the vector of ones, μ = population mean, Z = the known design matrix for genotypes, u = random genetic effects and 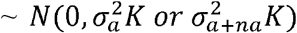 with K = kinship matrix, *a* = additive model, *na* = non-additive model, e = vector of residuals 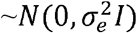.

**Table 3.** Proposed design matrix coding for auto-hexaploid sweetpotato as adapted from Slater et al. 2016.

|  | Pseudo_2x | Add_6x | Add+Non_6x |  |  |  |  |  |  |
| --- | --- | --- | --- | --- | --- | --- | --- | --- | --- |
| Effects/Marker | 1 | 1 | 1 | 2 | 3 | 4 | 5 | 6 | 7 |
| AAAAAA | 0 | 0 | 1 | 0 | 0 | 0 | 0 | 0 | 0 |
| AAAAAB | 1 | 1 | 0 | 1 | 0 | 0 | 0 | 0 | 0 |
| AAAABB | 1 | 2 | 0 | 0 | 1 | 0 | 0 | 0 | 0 |
| AAABBB | 1 | 3 | 0 | 0 | 0 | 1 | 0 | 0 | 0 |
| AABBBB | 1 | 4 | 0 | 0 | 0 | 0 | 1 | 0 | 0 |
| ABBBBB | 1 | 5 | 0 | 0 | 0 | 0 | 0 | 1 | 0 |
| BBBBBB | 2 | 6 | 0 | 0 | 0 | 0 | 0 | 0 | 1 |

Cross-validation was done by setting 20% of the population to missing phenotypes to be used as a validation set. We used 1,000 iterations to estimate the predictive ability of the models using both simple/oligo (quality traits in sweetpotato, disease traits in potato) and complex (storage root or tuber yield and yield component traits in both), as defined in Table 1.

Unlike in sweetpotato where phenotype and genotype data were balanced across experiments, (292 + Parents for DArTSeq and 315 + parents for GBSpoly), the potato experiments were unbalanced in terms of experimental genotypes. For the purposes of this study, we only selected the locations with the highest training population per trait. Consequently, we used AYP from Kunming (China; AYP_K), WMT from Kunming (China; WMT_K), LB from Oxapampa (Peru; LB2014_O), LB from Yunnan (China; LB2016_Y), PVY from Lima (Peru; PVY_L), TTW averaged across three treatments of 2016 in Ica (Peru; TTW16_Ica), and TTW in 2016 from Heilongjiang (China; TTW16_HLJ), all having number of genotypes indicated in Table 2. Differences in PA among models per trait were tested using t-tests. Quantitative-genetic parameters were tested for the additive model with or without dosage by obtaining the additive genetic variation 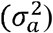 and random residual effects 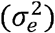 from the mixed linear model and calculating narrow-sense heritability for each trait as:

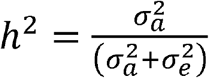

Additionally, we calculated the estimated rate of genetic gains from genomic selection per additive model with or without dosage for each trait according to **Oliveira et al. (2019)** as:

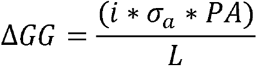

assuming L=5 for sweetpotato following the accelerated breeding scheme currently implemented (**Mwanga et al. 2017**), and L= 8 for potato.

### How many markers are adequate for prediction?

For sweetpotato, we used the GBSpoly data, using different filtration criteria to end up with different number of markers. We used three criteria i) total number of SNPs filtered at 10% MAF and ≥ 90% call rate, ii) total number of SNPs filtered at 30% MAF and ≥ 90% call rate (used in the analyses above), and iii) A random sample of 15,000 SNPs from the total number of SNPs and filtered at 30% MAF and ≥ 90% call rate. In Potato, the total number of SNPs was filtered using two criteria: i) 30% MAF and ≥ 90% call rate, ii) 40% MAF and ≥ 90% call rate. The model considering only additive effects was used in comparing the effect of number of markers in sweetpotato, while all three models were tested between the two filtering criteria in potato.

### Incorporating haplotypic-QTL in prediction models for sweetpotato

By taking advantage of the fully phased integrated linkage map from BT (**Mollinari et al. 2020**), we tested the predictive ability from QTL-informed models. Towards this end, we used the same cross-validation scheme as above, where 80%:20% random samples were used as training and testing populations, respectively, replicated 1,000 times. In order to detect QTL, we ran our random-effect multiple interval mapping (REMIM) using a sequential forward search (**Pereira et al. 2019**). We used score statistics to test map positions every 2 centiMorgans (cM) and added a QTL at a time using a relaxed genome-wide significance level threshold (α = 0.20). A window size of 20 cM was used to avoid that another position was selected very close to another QTL already in the model. For G-BLUP models, realized kinship matrices were based on the haplotype information from markers positioned every 2 cM in the genetic map. For QTL-BLUP (Q-BLUP), realized kinship matrices were based on the haplotypes from QTL peak marker; if there were more than one QTL, their kinship matrices were averaged out; if there were no QTL, we obtained the prediction as in G-BLUP. For Q+G-BLUP models, two terms were fitted, each with realized kinship matrices based on QTL peak markers (like for Q-BLUP) and the remaining markers in the linkage map but those selected as QTL.

## Results

### SNP profiles from the genotyping platforms

DArTseq sequencing of sweetpotato resulted in 13,504 biallelic SNPs (**Online Resource 2**). The call rates and polymorphic information content (PIC) are shown in Fig. 1A&B and ranged from about 0.4 - 1.0, with a mean of 0.96 for call rate and from 0 - 0.5 with a mean of 0.37 for PIC. Stringent filtering at a call rate ≥ 0.8 and PIC ≥ 0.25 left 9,649 SNPs that were used in AGHMatrix. Additional filtration in AGHMatrix at ≤ 0.1 missingness and ≥ 30% MAF resulted in 6,015 diploidized, biallelic SNPs being used to develop the matrices following additive (Add_2x_DArTSeq) and nonadditive (NonAdd_2x_DArTSeq) models.

**Fig. 1.**
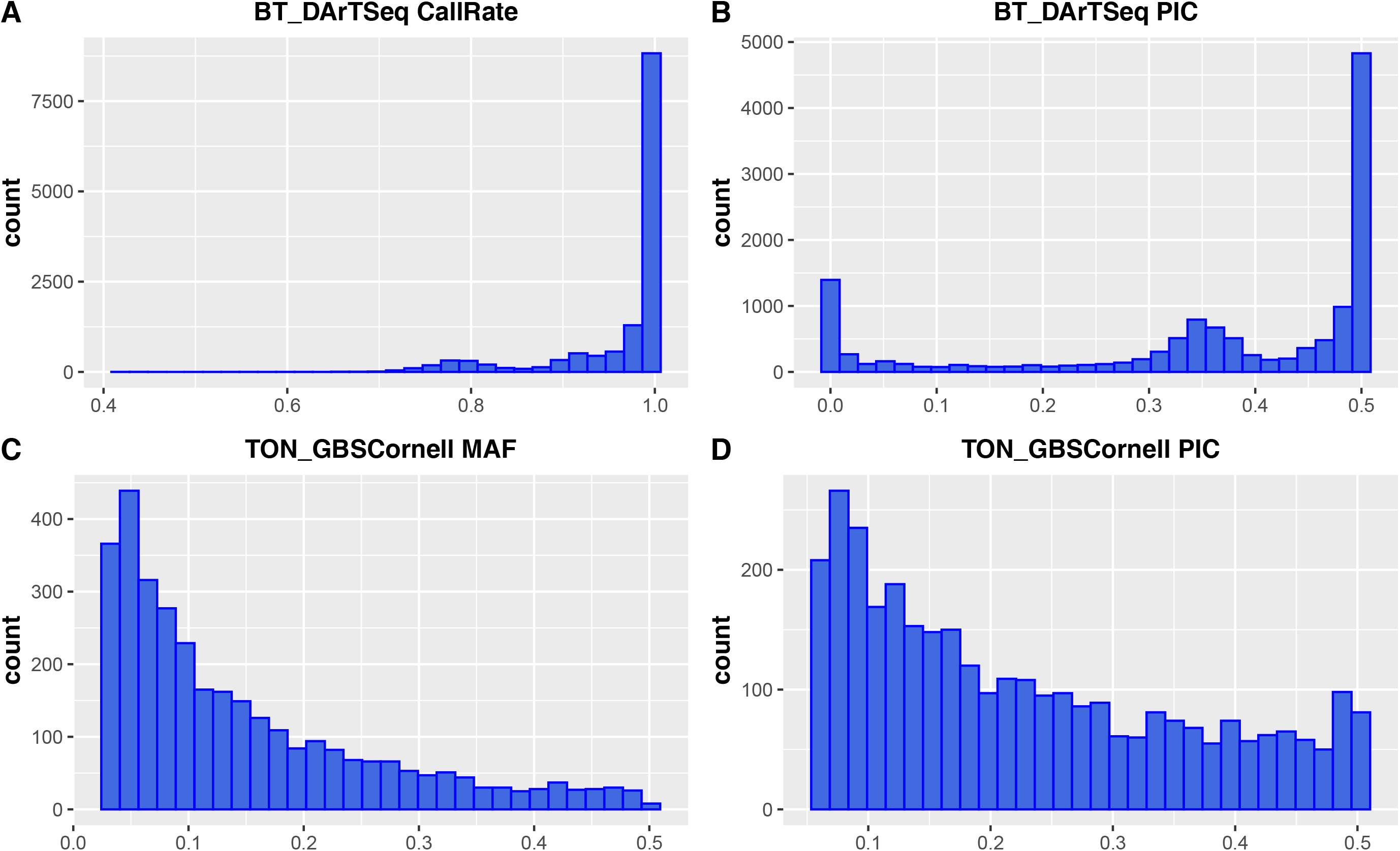
Quality attributes of the SNP profiles from DArTSeq (call rate and polymorphic information content; PIC) data in sweetpotato and GBSCornell (minor allele frequency; MAF and PIC) in potato

Cornell GBS in potato resulted in 295,401 biallelic SNPs at the variant calling step that were then hard-filtered to 3,262 high confidence SNPs by setting MAF ≥ 0.03, missing ≤ 2% and average read depth (DP) ≥ 16 (**Online Resource 3**). The 3,262 SNP profiles are shown in Fig. 1C&D showing MAF ranging from 0.03 - 0.5, with a mean of 0.15 and PIC ranging from 0.0 - 0.5, with a mean of 0.23. The 3,262 SNPs were used in the AGHMatrix relationship matrix development. For a relative comparison of models across crops for trait groups, we also filtered the Cornell GBS data in AGHMatrix at ≤ 0.1 missingness and ≥ 30% MAF as done for DArTSeq data above, which resulted in 411 SNPs used to develop the additive (Add_4x_GBSCornell), additive plus non-additive (Add+Non_4x_GBSCornell) and the pseudo-diploidized (Pseudo_2x_GBSCornell) models. Examining the relationship matrices indicated that at MAF ≥ 0.3, the full model (Add+Non_4x_GBSCornell) was mainly monomorphic. For potato therefore, we also changed the MAF to ≥ 0.4, which resulted in 178 SNPs that were used to develop a second set of relationship matrices. All PA comparisons among traits for potato are based on this matrix.

For GBSpoly in sweetpotato called according to **Wadl et al (2018)**, comparing diploid genotype data from M9 x M19 diploid population and hexaploid BT data showed that for the same level of genotype quality as for the diploid at about 25x depth of coverage, we needed ≥ 100x depth of coverage (Fig. 2). Consequently, for sweetpotato, GBSpoly data was filtered to this high depth of coverage, with MAF ≥ 0.05. This resulted in 34,390 high confidence SNPs (**Online Resource 4**) that were used in AGHMatrix to develop the additive (Add_6x_GBSpoly), additive plus non-additive (Add+Non_6x_GBSpoly) and the pseudo-diploidized (Pseudo_2x_GBSpoly) relationship matrices. The filters in AGHMatrix were set to ≤ 0.1 missing and ≥ 0.3 MAF as for the preceding data types and resulted in a final 2,883 SNPs that developed the matrices for model comparison. For comparing the effects of number of markers on PA, the first filtration criteria of 10% MAF and ≥ 90% call rate resulted in 10,358 SNPs, while the third criteria based on a random sample of 15,000 SNPs resulted in 1,291 SNPs that were used in PA comparison, based on the additive only model.

**Fig. 2.**
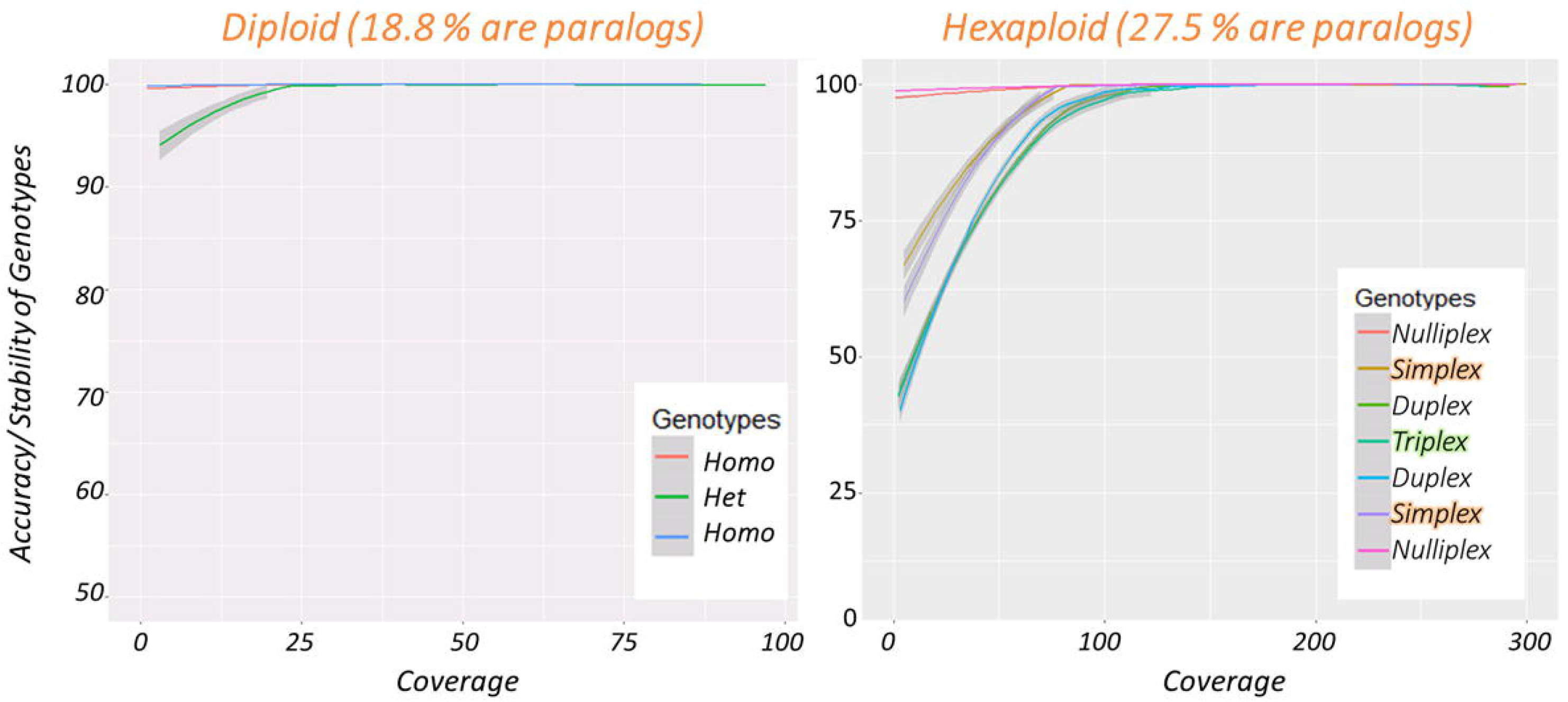
Comparison of genotype quality at different allele sequencing depths in diplod *I. trifida* (M9xM19) and hexaploid sweetpotato (*I. batatas*; BT)

The comparison of models with(out) QTL and use of markers *per se* or haplotypes was carried out using 30,684 SNPs from the same genotyping platform and data set, filtered and processed as described by **Mollinari et al. (2020)**, which were used to develop a 2,708.4 cM phased genetic linkage map for sweetpotato, and subsequent QTL analyses (**Pereira et al. 2019; Gemenet et al. 2019**). Sweetpotato BLUEs are provided in **Online Resource 5** while potato BLUEs are provided as **Online Resource 6**.

### Genotyping platforms, genetic effects and predictive ability

In sweetpotato, the diploidized additive model (Add_2x_DArTSeq) using data from DArTSeq performed equally highly as or sometimes better than the additive model using high confidence dosage data from GBSpoly (Add_6x_GBSpoly), depending on trait architecture, for simpler quality-related traits (Fig. 4). DM had 0.33 and 0.44, Starch had 0.32 and 0.38, BC had 0.43 and 0.43, FC_P had 0.44 and 0.45, while FC_U had 0.41 and 0.38 average PA for Add_2x_DArTSeq and Add_6x_GBSpoly models, respectively (Table 4). For these traits, additive only models were the best and the full model (Add_Non_6x_GBSpoly) always had negative PA due to a largely monomorphic relationship matrix. Nonetheless, the situation changed with yield-related traits as the effects of dosage and non-additive effects became more important. For these traits, the high-quality data with dosage from GBSpoly (Add_6x_GBSpoly) was always better in prediction when compared to the additive model with diploidized data (Add_2x_DArTSeq). NOCR had 0.19 and 0.31, TNR had 0.25 and 0.37, CYTHA had 0.18 and 0.22, RYTHA had 0.18 and 0.23, FYTHA had 0.21 and 0.26 average PA for Add_2x_DArTSeq and Add_6x_GBSpoly additive-models, respectively (Table 4). However, the additive only model with dosage (Add_6x_GBSpoly) was not always the best in PA for all yield-related traits, especially not for storage roots traits CYTHA and RYTHA, where it performed similar to either or both of the models considering non-additive effects whether with dosage (Add+Non_6x_GBSpoly) or without dosage (NonAdd_2x_DArTseq) (Fig. 3). Nevertheless, the largely monomorphic relationship matrix from the full model (Add+Non_6x_GBSpoly) ensured low predictive ability using this model for most yield-related traits as well, especially FYTHA which had the highest negative PA, (collapsed to zero in Fig.3, for plotting purposes). In general, pseudo-diploidizing data already called with dosage (Pseudo_2x_GBSpoly) drastically reduced PA even more than using data called as diploid (DArTseq). In potato, the situation was not very different as the diploidized additive model (Pseudo_2x_GBSCornell) was the second-best model after the additive only model with dosage (Add_4x_GBSCornell) for simpler disease traits and its comparative advantage significantly reduced with more complex traits (Fig. 4). LB2014_O had 0.68 and 0.63, LB2016_Y had 0.62 and 0.52, PVY_L had 0.54 and 0.50, AYP_K had 0.45 and 0.34, WMT_K had 0.48 and 0.34, TTW16_Ica had 0.16 and 0.16, while TTW16_HLJ had 0.37 and 0.32 average PA for Add_4x_GBSCornell and Pseudo_2x_GBSCornell, respectively. As with the full model in sweetpotato, the model including non-additive effects (Add+Non_4x_GBSCornell) was the least performing in terms of PA (Table 4).

**Table 4.** Summary quantitative-genetic parameters derived from genomic selection with cross validation applying different genetic effects models in sweetpotato and potato. 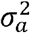 is the additive genetic variation, 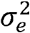 the residual variance, *h*^2^ the narrow-sense heritability, *PA* the predictive ability, ∆*GG* the estimated rate of genetic gain considering the current breeding cycle length

| Crop | Trait<sup>a</sup> | Model<sup>b</sup> | $\sigma_a^2$ | $\sigma_e^2$ | $h^2$ | $PA$ | $\Delta GG$ |
| --- | --- | --- | --- | --- | --- | --- | --- |
| Sweetpotato | DM | Add_2x_DArTSeq | 1.6935 | 2.9536 | 0.36 | 0.33b | 0.085889 |
|  |  | Add_6x_GBSpoly | 4.0035 | 2.0762 | 0.66 | 0.44a | 0.176077 |
|  | Starch | Add_2x_DArTSeq | 6.1716 | 13.7616 | 0.31 | 0.32b | 0.158993 |
|  |  | Add_6x_GBSpoly | 12.1683 | 11.5424 | 0.53 | 0.38a | 0.265111 |
|  | BC | Add_2x_DArTSeq | 150.2336 | 113.1697 | 0.57 | 0.43a | 1.0541 |
|  |  | Add_6x_GBSpoly | 225.1431 | 152.0581 | 0.60 | 0.43a | 1.29041 |
|  | FC_P | Add_2x_DArTSeq | 0.5416 | 0.3304 | 0.62 | 0.44a | 0.064762 |
|  |  | Add_6x_GBSpoly | 0.8168 | 0.4189 | 0.66 | 0.45a | 0.081339 |
|  | FC_U | Add_2x_DArTSeq | 12.9633 | 10.9257 | 0.54 | 0.41a | 0.295238 |
|  |  | Add_6x_GBSpoly | 16.1489 | 16.777 | 0.49 | 0.38ab | 0.305411 |
|  | NOCR | Add_2x_DArTSeq | 50134102 | 2.93E+08 | 0.15 | 0.19b | 269.0607 |
|  |  | Add_6x_GBSpoly | 1.36E+08 | 2.43E+08 | 0.36 | 0.31a | 722.4699 |
|  | TNR | Add_2x_DArTSeq | 1.86E+08 | 7.40E+08 | 0.20 | 0.25b | 681.9091 |
|  |  | Add_6x_GBSpoly | 4.71E+08 | 5.83E+08 | 0.45 | 0.37a | 1606.149 |
|  | CYTHA | Add_2x_DArTSeq | 8.6149 | 27.7157 | 0.13 | 0.18b | 0.105664 |
|  |  | Add_6x_GBSpoly | 8.6149 | 26.6061 | 0.24 | 0.22a | 0.129145 |
|  | RYTHA | Add_2x_DArTSeq | 4.6249 | 31.4849 | 0.13 | 0.18b | 0.07742 |
|  |  | Add_6x_GBSpoly | 10.9811 | 29.4989 | 0.27 | 0.23a | 0.152434 |
|  | FYTHA | Add_2x_DArTSeq | 7.678 | 26.0083 | 0.23 | 0.21b | 0.116379 |
|  |  | Add_6x_GBSpoly | 12.8721 | 26.6023 | 0.33 | 0.26a | 0.186564 |
| Potato | LB2014_O | Add_4x_GBSCornell | 0.0189 | 0.0193 | 0.49 | 0.68a | 0.011686 |
|  |  | Pseudo_2x_GBSCornell | 0.0195 | 0.023 | 0.46 | 0.63b | 0.010997 |
|  | LB2016_Y | Add_4x_GBSCornell | 0.0191 | 0.0259 | 0.42 | 0.62a | 0.010711 |
|  |  | Pseudo_2x_GBSCornell | 0.0166 | 0.0323 | 0.34 | 0.52b | 0.008375 |
|  | PVY_L | Add_4x_GBSCornell | 0.0419 | 0.0738 | 0.36 | 0.54a | 0.013817 |
|  |  | Pseudo_2x_GBSCornell | 0.0364 | 0.0818 | 0.31 | 0.50ab | 0.011924 |
|  | AYP_K | Add_4x_GBSCornell | 0.0118 | 0.0327 | 0.27 | 0.45a | 0.00611 |
|  |  | Pseudo_2x_GBSCornell | 0.0066 | 0.0389 | 0.15 | 0.34b | 0.003453 |
|  | WMT_K | Add_4x_GBSCornell | 0.0132 | 0.0322 | 0.29 | 0.48a | 0.006893 |
|  |  | Pseudo_2x_GBSCornell | 0.0069 | 0.0392 | 0.15 | 0.34b | 0.00353 |
|  | TTW16_Ica | Add_4x_GBSCornell | 2.00E-04 | 0.0028 | 0.07 | 0.16a | 0.000283 |
|  |  | Pseudo_2x_GBSCornell | 3.00E-04 | 0.0027 | 0.10 | 0.16a | 0.000346 |
|  | TTW16_HLJ | Add_4x_GBSCornell | 0.0061 | 0.018 | 0.25 | 0.37a | 0.003612 |
|  |  | Pseudo_2x_GBSCornell | 0.0049 | 0.0192 | 0.20 | 0.32b | 0.0028 |

<sup>a</sup>Traits as defined in Table 1, <sup>b</sup>Models: Add\_2x\_DArTseq = additive model using data from DArTseq called as diploid; Add\_6x\_GBSpoly= additive model using data with dosage from GBSpoly; Add\_4x\_GBSCornell = additive
model using data with dosage from GBS at Cornell, Pseudo\_2x\_GBSCornell = additive model using data from GBS Cornell with three heterozygote classes collapsed into one.

**Fig. 3.**
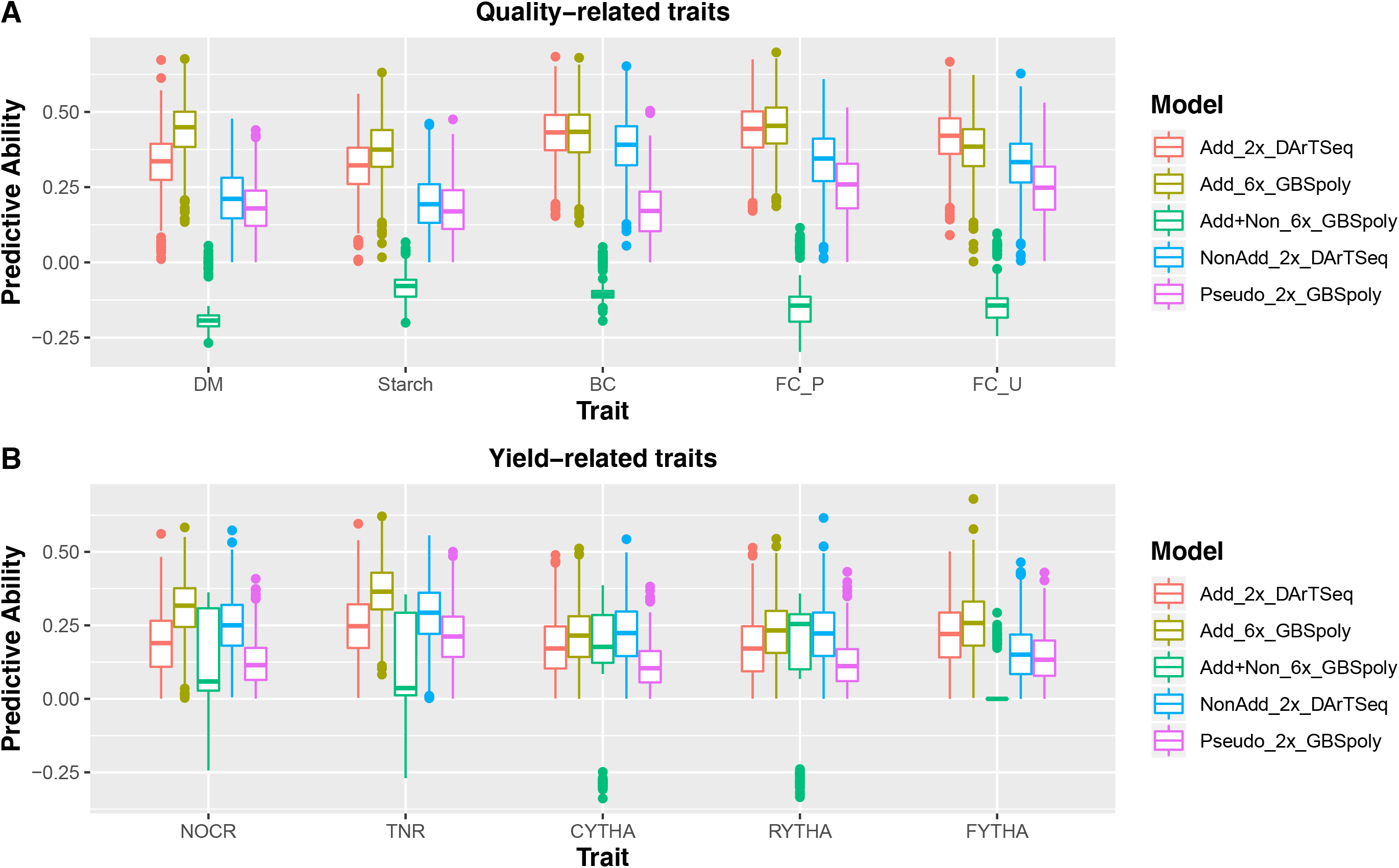
Boxplots comparing predictive ability of additive-effects-only models without dosage (Add_2x_DArTseq) and with dosage (Add_6x_GBSpoly); models considering also non-additive effects (NonAdd_2x_DArTSeq; Add+Non_6x_GBSpoly); and pseudo-diploidized dosage data (Pseudo_2x_GBSpoly) for quality related traits (A; DM = dry matter, starch, BC = β-carotene, FC_P = flesh color in Peru; FC_U = flesh color in Uganda); and yield related traits (B; NOCR = number of commercial storage roots, TNR = total number of storage roots, CYTHA = weight of commercial storage roots, RYTHA = weight of total storage roots, FYTHA = total weight of foliage) in a full-sib family of sweetpotato.

**Fig. 4.**
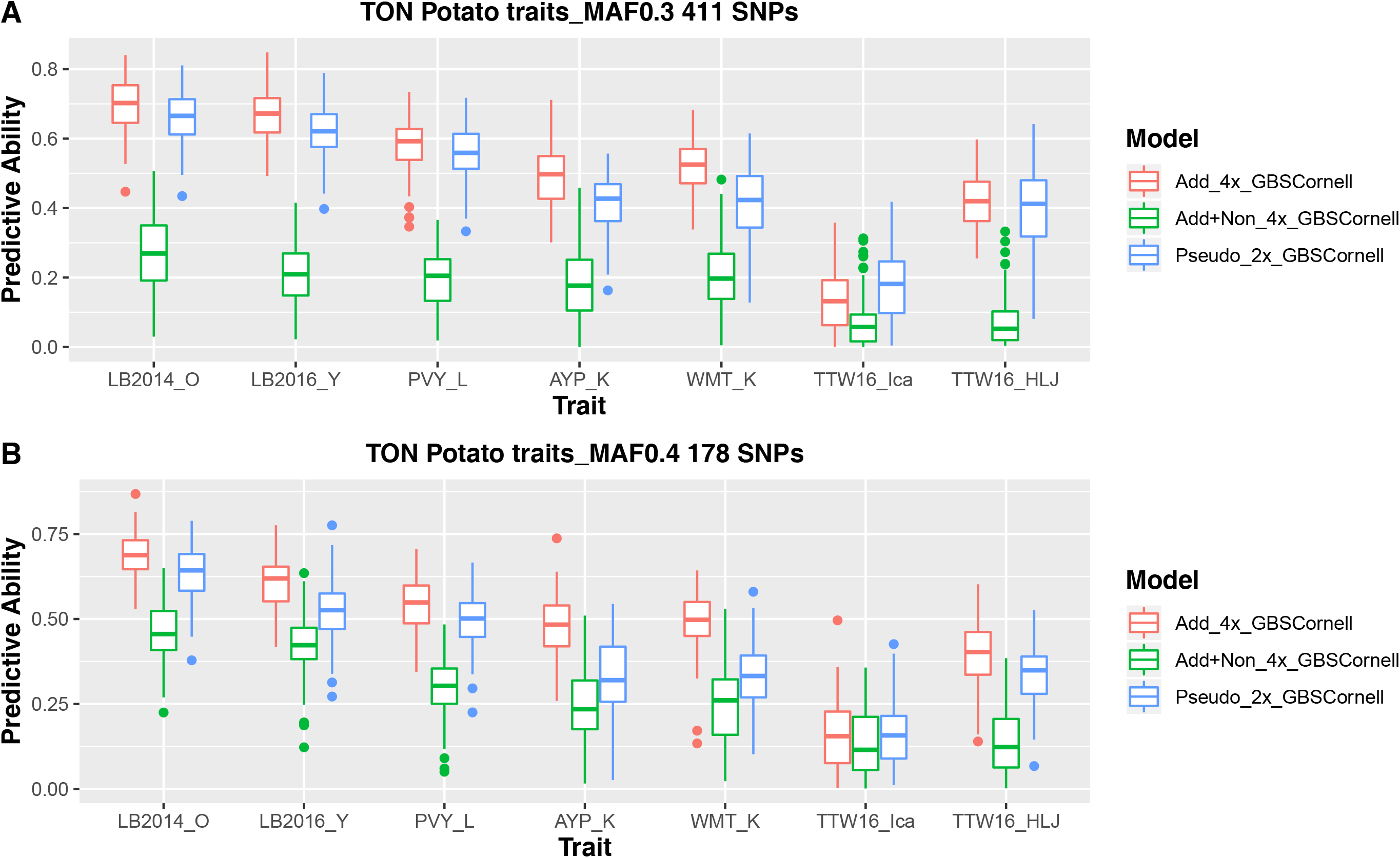
Box plots comparing predictive ability of additive-effects-only model (Add_4x_GBSCornell); additive and non-additive effects (Add+Non_4x_GBSCornell); and pseudo-diploidized dosage data (Pseudo_2x_GBSCornell); using minimum allele frequency (MAF) ≥ 30% (A; 411 SNPs) and MAF ≥ 40% (B; 178 SNPs). LB2014_O = late blight in Oxapampa (Peru) in 2014, LB2016_Y = late blight in Yunnan (China) in 2016, PVY_L = potato virus Y in Lima (Peru), AYP_K = average yield per plant in Kunming (China), WMT_K = weight of marketable tubers in Kunming, TTW16_Ica = total tuber weight in Ica (Peru) in 2016 across three drought treatments, TTW16_HLJ = total tuber weight in Heilongjiang (China) in 2016, single treatment, in potato.

### Number of markers and environments

Our results in potato indicated that an increased number of markers by more than double did not have a significant effect on PA (411 vs 178 SNPs; Fig. 4A) considering the best predictive model (Add_4x_GBSCornell). LB2014_O had 0.69 and 0.68, LB2016_Y had 0.66 and 0.62, PVY_L had 0.59 and 0.54, AYP_K had 0.51 and 0.45, WMT had 0.51 and 0.48, TTW16_Ica had 0.19 and 0.16, while TTW16_HLJ had 0.40 and 0.37 average PA for 411 and 178 SNPs respectively. Similarly, in sweetpotato, comparing PA using 10,358 SNPs, 2,883 SNPs and 1,291 SNPs using the best predictive model (Add_6x_GBSpoly) showed no effect of increasing marker density at the cost of marker informativeness on PA. PA based on 10,358 SNPs which had 10% MAF generally performed lower than 2,883 and 1,291 SNPs which both had 30% MAF (Fig. 5). Additionally, 2,883 SNPs did not have a clear comparative advantage over 1,291 SNPs (Fig. 5). Regarding traits in different locations, environmental effects on PA were observed, though the magnitude of such effects was also dependent on trait architecture. The PA based on the best model for FC_P (0.45; Peru) and FC_U (0.38; Uganda) in sweetpotato and LB2014_O (0.68; Peru) and LB2016_Y (0.62; China), in potato, though a bit different, were both relatively high to allow meaningful selections for the trait. For more complex yield traits, the PA for TTW16_Ica (0.16; Peru) and TTW16_HLJ (0.37; China) were significantly different.

**Fig. 5.**
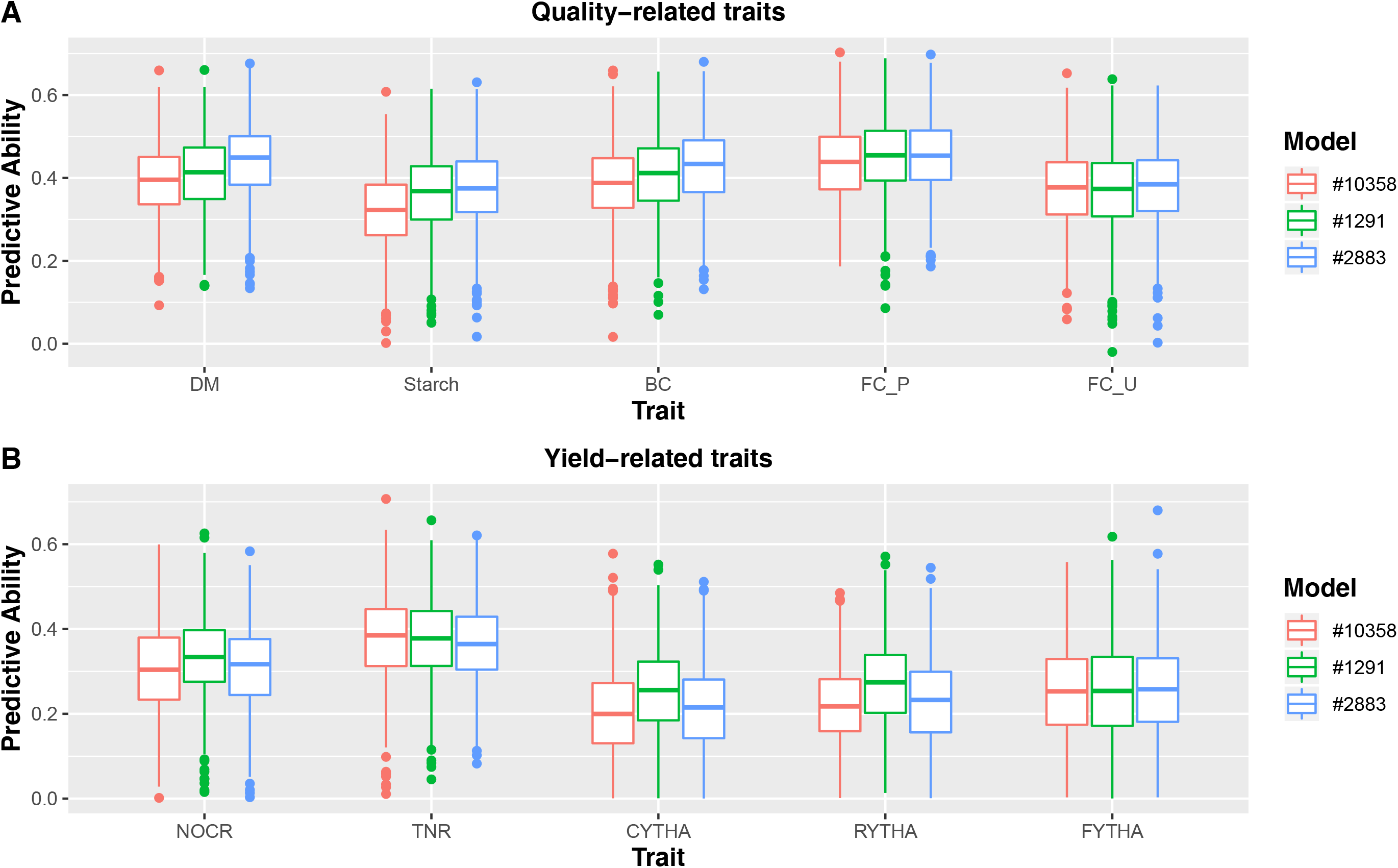
Box plots comparing the effect of number of markers on predictive ability using additive-effects only model (Add_6x_GBSpoly) with 10,358 SNPs, 2,883 SNPs and 1,291 SNPs in sweetpotato. A; DM = dry matter, starch, BC = β-carotene, FC_P = flesh color in Peru; FC_U = flesh color in Uganda; and yield related traits: B; NOCR = number of commercial storage roots, TNR = total number of storage roots, CYTHA = weight of commercial storage roots, RYTHA = weight of total storage roots, FYTHA = total weight of foliage in a full-sib family of sweetpotato.

### Effects of quantitative trait loci, haplotypes and dosage on predictive ability

We additionally tested three analysis models using BT sweetpotato data: i) Q-BLUP based on relationship matrices from QTL-peak haplotypes, ii) Q+G-BLUP fitting two terms based on QTL-peak haplotypes and the rest of the markers in the linkage map, iii) G-BLUP, predictions using markers spaced every 2 cM in the genetic map without considering QTL. The PA results are shown in Fig. 6. Considering QTL haplotypes either *per se* (Q-BLUP) or with G-BLUP (Q+G-BLUP) had a clear comparative advantage for PA in simpler traits. However, this comparative advantage faded with more complex yield-related traits. Our results therefore show that with genomic selection, the comparative advantage of using the linkage map information and QTL is dependent on trait architecture, hence the magnitude of QTL effects that can be mapped (Fig. 6).

**Fig. 6.**
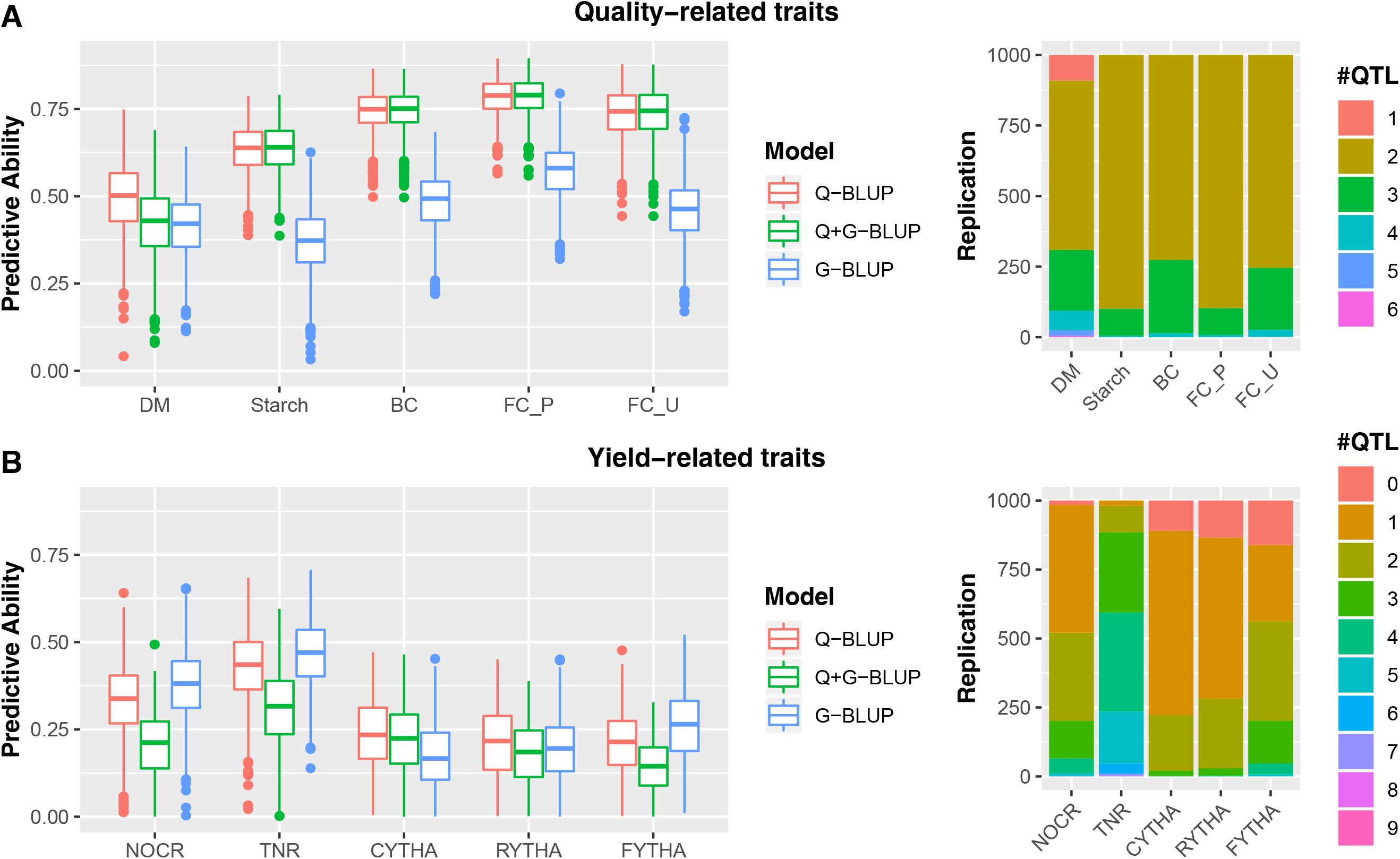
Boxplots comparing predictive ability of models using QTL haplotypes only in prediction (Q-BLUP); QTL combined with prediction based on markers per se, (Q+G-BLUP); prediction using markers per se without QTL (G-BLUP) for quality-related traits (A; DM = dry matter, starch, BC = β-caroten, FC_P = flesh color in Peru; FC_U = flesh color in Uganda); and yield related traits (B; NOCR = number of commercial storage roots, TNR = total number of storage roots, CYTHA = weight of commercial storage roots, RYTHA = weight of total storage roots, FYTHA = total weight of foliage) in a full-sib family of sweetpotato.

### Genetic variation, heritability and estimated rate of genetic gain

Given that the additive effects only model with dosage performed better for most traits in both sweetpotato and potato (Add_6x_GBSpoly and Add_4x_GBSCornell, respectively), we evaluated quantitative genetic parameters for this model in comparison with the additive model without dosage for both crops (Add_2x_DArTseq for sweetpotato and Pseudo_6x_for potato). Narrow sense heritability (h^2^) ranged from 0.24-0.66 for the model with dosage (Add_6x_GBSpoly) and 0.13-0.62 for the model without dosage (Add_2x_DArTseq) in sweetpotato. In potato, (h^2^) ranged from 0.07 – 0.49 in the model with dosage (Add_4x_GBSCornell) and 0.10 – 0.46 in the model with pseudo-diploidized dosages (Pseudo-2x_GBSCornell; Table 4). As expected, traits with simpler architecture (quality-related traits in sweetpotato; disease traits in potato had the highest (h^2^) compared to more complex yield-related traits. All models across crops resulted in positive estimated genetic gain considering L= 5 years in sweetpotato and L= 8 in potato, which are the cycle lengths of current breeding schemes at CIP (Table 4). This implies that more genetic gains can be realized if such breeding cycle lengths are further significantly reduced.

## Discussion

### Low-cost, targeted amplicon sequencing platforms could realize faster genetic gains per unit time

Having a reliable, cost-efficient genotyping platform that ensures faster data turn-around to breeding programs on time to impact selection and advancement decisions is a must for routine application of genomic selection in plant breeding programs. Here we have compared results based on data from three GBS-based platforms, two of which provide data at the commercial diploid sequencing depth level (DArTSeq and GBS-Cornell). About 100x read depth was required to confidently call all the five heterozygous dosage classes of sweetpotato, against 25-30x required for the diploid. These results agree with studies in potato where **Uitdewilligen et al. (2013)** reported that 60-80x depth was required to confidently call the three heterozygote classes. GBSpoly (**Wadl et al. 2018**) which had high quality dosage data in our study was developed as part of a project to understand optimal conditions for GBS in hexaploid sweetpotato and therefore not amenable to routine use in plant breeding. Other options for more precise genotyping such as SNP arrays, in addition to issues with ascertainment biases, are crop-specific and therefore do not benefit from economies of scale that drive costs down. Breeding programs of polyploid crops therefore have to weigh whether investing more for higher depth of sequencing is an efficient resource allocation strategy (**Endelman et al. 2018**). To this end, although our results show that genotype quality and consequently the number of realized SNPs is lower with low allele sequencing depth, we also show as described in the next sections that only a small number of highly informative SNPs are required to realize relatively high PA depending on the trait. These results agree with the findings of **Chang et al. (2019)** who showed that PA can be improved by prioritizing relevant SNP polymorphisms. This therefore implies that for practical plant breeding applications, using established genotyping platforms that ensure low-costs due to scale effects and faster data turn-around will have better likelihood of success in routine application of genomic selection in polyploids despite the low allele sequencing depths. Since both crops already have GBS-based SNPs at high density, the process can be fast-tracked by targeting the high informative segregating loci in amplicon sequencing. This is encouraging as polyploid crops in developing countries with limited access to expensive, high quality genotypic datasets could also deploy GS approaches.

### A few highly informative SNPs segregating in the population are adequate for prediction purposes

**Guo et al. (2018)** found that at allele sequencing depth between 10x to 20x, between 80-100K SNPs would be required to accurately predict additive breeding values in tetraploid ryegrass. Our research in both hexaploid sweetpotato and tetraploid potato however shows a reduced number of realized SNPs after quality filtration, which can be attributed to the difficulty of genotyping polyploid crops. SNP calling in polyploids is further complicated by the presence of polymorphic positions across homologues within and among individuals in addition to the polymorphic positions within a single homologue among individuals (**Clevenger et al. 2015**). In our potato example, the initial filtration of SNPs to allele sequencing depth at DP ≥ 16 and MAF 3% resulted in only 3,262 SNPs. The same scenario was observed for sweetpotato. However, our results also show that if SNPs are highly informative (MAF ≥ 30%), a number as low as 178 SNPs could give relatively high PA comparable to a larger number of SNPs. In potato, 178 SNPs at MAF ≥ 40% performed relatively similar as 411 SNPs at MAF ≥ 30%. Not shown results from a preliminary analysis of the same dataset of potato using MAF ≥ 10% resulted in 1,710 SNPs whose PA did not differ significantly with the PA using either 411 or 178 SNPs. Additionally, in sweetpotato, 2,883 SNPs at MAF ≥ 30% gave the same or better PA as 10,358 SNPs at MAF ≥ 10%, and 1,291 SNPs. Our results therefore agree with the findings of **Covarrubias-Pazaran et al. (2018)** using three biparental populations of the American cranberry, that addition of SNPs after 500 markers did not result in much increase in PA as only a few hundred SNPs were needed to reach PA plateau. Even though their study used a consensus map to intentionally distribute markers evenly across the genome, our random sampling method based on MAF and PIC came to the same conclusion. These results imply that breeding programs with limited resources for genotyping can target few highly informative regions within the genome that are segregating in their breeding populations via targeted genotyping methods following amplicon sequencing techniques, as a cost-effective way of incorporating genomic selection in their breeding programs. We propose the use of between 500-1000 highly informative SNPs for routine prediction purposes in a breeding program.

### Modelling non-additive genetic effects has negligible contribution to predictive ability

Our results both in potato and sweetpotato show that additive effects-only models, whether diploidized or with dosage, were comparatively better in PA than the models considering non-additive effects for all simple traits. This comparative advantage however lessened with more complex traits, where non-additive effects and inclusion of dosage information became slightly more relevant, although in most cases the additive effects-only model with dosage still remained the best in terms of PA. This finding makes sense in quantitative genetic terms as the more the number of genes affecting a trait, the more the expected interaction among loci. In sweetpotato for example, issues of ‘missing’ heritability have been established for yield-related traits using the current BT population in multiple environments, where only a few QTL with very small effects were reported even though a very dense, well phased hexaploid genetic map was used (**Pereira et al. 2019; Gemenet et al. 2020**). According to **Varona et al. (2018)**, the contribution of non-additive effects to genetic variance depends on the allele frequency of the causative loci, and their consideration in breeding programs can improve the prediction accuracy for breeding values and inform cross-combinations that maximize non-additive variation in progeny. Several studies have however shown that inclusion of non-additive effects in the prediction models have negligible effects in improving the accuracy of predicting breeding (additive) values. For instance, **Endelman et al. (2018)** reported uncertainty in partitioning non-additive genetic variance in tetraploid potato, whereas **Crow (2010)**, suggested that variance due to epistasis would have little effects in plant breeding as additive variance and covariance effects quickly overshadow such contribution following selection. Non-additive effects are mainly considered important in genomic prediction (prediction for performance of different traits based on the genotype of the individual), while additive-only methods as important in genomic selection (prediction of parental value of an individual), because only additive effects can be passed from parents to progeny (**Varona et al. 2018**). However, our results, supported by previous findings in other crops, imply that in light of the large number of moving parts to consider, including concerns with genotyping platforms and genotype quality for polyploids, practical breeding programs for potato and sweetpotato, and perhaps other polyploid crops, will achieve more advances considering only the infinitesimal model (additive) for both genomic selection and genomic prediction.

### The relative importance of considering dosage, haplotypes and quantitative trait loci is dependent on trait architecture

**Oliveira et al. (2019)** showed that the relative advantage of including dosage information to PA is dependent on trait architecture. Our results confirm this and show that for simple traits diploidized data, especially when the genotypic data are directly called as diploid during variant calling e.g. the DArTSeq data in sweetpotato rather than pseudo-diploidizing data already called with dosage e.g. in GBS-Cornell data in potato, would just do fine. However, as the traits become more complex, considering dosage improves PA and therefore the rate of progress that can be made for such traits. **Endelman et al. (2018)** also showed that not considering allele dosage effects in potato reduced prediction accuracy by about 0.13 on average using data from the SolCAP potato SNP array, where they reported PA ranging from 0.06 to 0.63 for specific gravity, yield and fry color. Given that most traits are quantitative, we recommend the use of data with dosage even though they may come from sequencing platforms with low allele sequencing depth, that could benefit more with improved genotype calling methods, such as Bayesian genotype calling methods.

Our data also shows that for all traits, considering both QTL and haplotypes resulted in the best PA especially for simple traits, although this comparative advantage also faded with more complex yield traits. Having markers in complete LD with causative QTL for a given trait is a prerequisite for improving PA in genomic prediction (**Velasco et al. 2019**). The study of **Cuyabano et al. (2014)** showed that considering haplotype blocks rather than single markers improved PA for dairy traits in cattle. This is because haplotypes are supposed to be in tighter LD with QTL than single markers. This can be attributed to the fact that GS-only GBLUP methods use the average genome information relationship for model building and for prediction whereas incorporating QTL analysis gives different weights (QTL effects) to different “significant” genome positions (QTL positions) for model building and for prediction. Due to this, studies have proposed a combination of QTL mapping to explain trait architecture and genomic prediction, to improve PA (**Spindel et al. 2016; Lopes et al. 2017; Morgante et al. 2018; Bhandari et al. 2019**). Our results however indicate that the relative advantage of considering QTL-based haplotypes is dependent on trait architecture and directly related to the number and effect size of the QTL in question. In this case, yield-related traits did not show much improvement in PA when QTL were considered. Despite this finding, additional efforts in studying the effect of haplotype structure on PA is recommended to increase the likelihood of fully recovering the polyploid genetic information, where the information from individual dosage markers can be rather limited. However, given that QTL mapping/GWAS methods require high density markers, the application of such a strategy should be considered in the context of the cost of developing high density markers against available resources for genotyping in a given program. Additionally, such methods would be computationally demanding and should also be considered depending on available computational tools and analytical capacity of a given program.

### Further considerations for optimized breeding programs using genomic selection

The PA of genomic selection is influenced by several factors including trait architecture, the size of the training population, the relationship between the training and validation populations, heritability of the trait, the level of linkage disequilibrium (LD), marker density, environmental variances and covariance among traits (**Nakaya and Isobe 2012**). In addition to the already discussed factors, our results indicate that environment plays a significant role in determining PA as can be seen in the same traits measured across several environments. Additionally, PA magnitude even for simple traits were lower in sweetpotato where we used BLUEs across six environments, than in potato where predictions were made per single environment. Models incorporating genotype-x-environment interaction are important and more realistic when predicting performance of untested genotypes across environments (**Burgueno et al. 2012; Heslot et al. 2014; Wang et al. 2018**). Furthermore, PA for complex yield-related traits were always lower than for simpler quality-related or disease traits. PA for such complex traits have been shown to benefit from multi-trait selection models incorporating simpler, correlated traits with the primary trait (**Covarrubias-Pazaran et al. 2018; Michel et al. 2019).** Additionally, **Bernal-Vasquez et al. (2014)** alluded to the fact that phenotypic data analysis contributed to improved PA, which speaks to the necessary precision and accuracy of the phenotype in training populations. Taken together, the current results show that genomic selection will contribute towards increased genetic gains, especially via reduced breeding cycle time in potato and sweetpotato. However, the effectiveness of genomic selection will have to be considered from the perspective of optimizing the entire breeding program (**Cobb et al. 2019**). This refers to the assembly and deployment of a package of technological tools that allow a specific program to realize maximum genetic gains within its current context in terms of time and resources, by exploiting all components of the breeder’s equation. Therefore, given the diversity existing from program to program in terms of resources and human capacity, no ‘one size fits all’ scenario is anticipated.

Finally, it does not escape to our attention that the predictions here-in are based on single populations. However, plant breeding requires several levels of allele recombination through generations. We cannot estimate from the current data, how such recombination complexity will affect the efficiency of GS in breeding programs. Additional studies estimating PA in actual multigeneration breeding populations therefore need to be carried out to reliably estimate the value of GS to potato and sweetpotato, and perhaps other polyploid breeding programs.

## Supporting information

Online Resource 1

Online Resource 2

Online Resource 3

Online Resource 4

Online Resource 5

Online Resource 6

## Data availability

All single nucleotide polymorphism (SNP) data used in the current manuscript are provided with the manuscript as **Online Resources 2-4** while all best linear unbiased estimators (BLUEs) are provided as **Online Resources 5 and 6**.

## Acknowledgements

The sweetpotato research, most co-authors and GBSpoly genotyping was supported by a Bill & Melinda Gates Foundation grant (Grant number OPP1052983). DArTSeq genotyping of sweetpotato was done through the collaboration of the Integrated Genotyping Service and Support platform (IGSS) and DArT^TM^ in Australia. Potato research and genotyping at Cornell was supported by a GIZ grant for the project ‘Accelerating the Development of Early-Maturing-Agile Potato for Food Security through a Trait Observation and Discovery Network’. The work was done as part of the Consultative Group on International Agricultural Research (CGIAR)-Research Program on Roots, Tubers and Bananas (RTB) which is supported by CGIAR Fund Donors (http://www.cgiar.org/about-us/our-funders/). Research teams at CIP and collaborating partners in China are greatly appreciated for providing phenotypic data.

## Online Resource Captions

**Online Resource 1** Variant calling pipeline used in the GBSapp for calling GBSpoly data in sweetpotato

**Online Resource 2** DArTSeq SNP data for the Beauregard x Tanzania (BT) sweetpotato full-sib family

**Online Resource 3** GBS-Cornell SNP data for the trait observation network (TON) potato population

**Online Resource 4** GBSpoly SNP data for the Beauregard x Tanzania (BT) sweetpotato full-sib family

**Online Resource 5** Best linear unbiased estimators for sweetpotato traits used in genomic prediction in the current study

**Online Resource 6** Best linear unbiased estimators for potato traits used in genomic prediction in the current study

## References

Altshuler D, Pollara VJ, Cowles CR, Van Etten WJ, Baldwin J, et al (2000) An SNP map of the human genome generated by reduced representation shotgun sequencing. Nature 407: 513–516

Amadeu RR, Cellon C, Olmstead JW, Garcia AA, Resende MF, Muñoz PR (2016) AGHmatrix: R package to construct relationship matrices for autotetraploid and diploid species: A blueberry example. Plant Genome doi: 10.3835/plantgenome2016.01.0009

Baird NA, Etter PD, Atwood TS, Currey MC, Shiver AL, et al (2008) Rapid SNP discovery and genetic mapping using sequenced RAD markers. PLoS One 3: e3376

Bernal-Vasquez A-M, Möhring J, Schmidt M, Schönleben M, Schön C-C, Piepho H-P (2014) The importance of phenotypic data analysis for genomic prediction - a case study comparing different spatial models in rye. BMC Genomics 15:646

Bhandari A, Bartholome J, Cao-Hamadoun T-V, Kumari N, Frouin J, Kumar A, et al (2019) Selection of trait-specific markers and multi-environment models improve genomic predictive ability in rice. PLoS ONE 14(5): e0208871

Blischak PD, Kubatko LS, Wolfe AD (2016) Accounting for genotype uncertainty in the estimation of allele frequencies in autopolyploids. Mol Ecol Resources 16:742–754

Blischak PD, Kubatko LS, Wolfe AD (2018) SNP genotyping and parameter estimation in polyploids using low-coverage sequencing data. Bioinformatics 34(3):407–415

Burgueño J, de los Campos G, Weigel K, Crossa J (2012) Genomic prediction of breeding values when modeling genotype x environment interaction using pedigree and dense molecular markers. Crop Sci 52:707–719

Chang L-Y, Toghiani S, Aggrey SE, Rekaya R (2019) Increasing accuracy of genomic selection in presence of high-density marker panels through the prioritization of relevant polymorphisms. BMC Genetics 20:21

Clark SA, van der Werf J (2013) Genomic best linear unbiased prediction (gBLUP) for the estimation of breeding values. Methods Mol Biol 1019:321–330

Clevenger J, Chavarro C, Pearl SA, Ozias-Akins P, Jackson SA (2015) Single nucleotide polymorphism identification in polyploids: A review, example, and recommendations. Mol Plant 8:831–846

Cobb JN, Juma RU, Biswas PS, Arbelaez JD, Rutkoski J, Atlin G, Hagen T, Quinn M, Ng EH (2019) Enhancing the rate of genetic gain in public-sector plant breeding programs: lessons from the breeder’s equation. Theor Appl Genet 132-627–645

Courtois B, Audebert A, Dardou A, Roques S, Ghneim-Herrera T, et al (2013) Genome-wide association mapping of root traits in a japonica rice panel. PLoS ONE 8(11): e78037

Covarrubias-Pazaran G, Schlautman B, Diaz-Garcia L, Grygleski E, Polashock J, et al (2018) Multivariate GBLUP improves accuracy of genomic selection for yield and fruit weight in biparental populations of Vaccinium macrocarpon Ait. Front. Plant Sci. 9: 1310

Crow, J. F. (2010). On epistasis: why it is unimportant in polygenic directional selection. Philos. Trans. R. Soc. Lond. B Biol. Sci. 365, 1241–1244

Cruz VM, Kilian A, Dierig DA (2013) Development of DArT marker platforms and genetic diversity assessment of the U.S. collection of the new oilseed crop lesquerella and related species. PLoS One 8(5): e64062

Cuyabano BCD, Su G Lund MS (2014) Geneomic prediction of genetic merit using LD-based haplotypes in the Nordic Holstein population. BMC Genomics 15:1171

Elshire RJ, Glaubitz JC, Sun Q, Poland JA, Kawamoto K, et al (2011) A robust, simple genotyping-by-sequencing (GBS) approach for high diversity species. PLoS One, 6(5):e19379

Endelman JB, Carley CAS, Bethke BC, Coombs JJ, Clough ME et al (2018). Genetic variance partitioning and genome-wide prediction with allele dosage information in tetraploid potato. Genetics 209:77–87

Faville MJ, et al (2018) Predictive ability of genomic selection models in a multi-population perennial ryegrass training set using genotyping-by-sequencing. Theor Appl Genet 131:703–720

Gemenet DC, Pereira GDS, De Boeck B, Wood JC, Mollinari M, Olukolu BA et al (2020) Quantitative trait loci and differential gene expression analyses reveal the genetic basis for negatively-associated β-carotene and starch content in hexaploid sweetpotato [Ipomoea batatas (L.) Lam.]. Theor App Genet 133:23–36

Grüneberg W, Mwanga R, Andrade M, Espinoza J (2009) Breeding clonally propagated crops. In FAO, selection methods: chapter 13, part 5

Guo X, Cericola F, Fè D, Pedersen MG, Lenk I, Jensen CS, Jensen J and Janss LL (2018) Genomic prediction in tetraploid ryegrass using allele frequencies based on genotyping by sequencing. Front Plant Sci 9:1165

Heslot N, Akdemir D, Sorrells ME, Jannink JL (2014) Integrating environmental covariates and crop modeling into the genomic selection framework to predict genotype by environment interactions. Theor Appl Genet 127:463–480

Kilian A et al (2012) Diversity Arrays Technology: A generic genome profiling technology on open platforms. Methods Mol Biol 888:67–89

Lipka AE, Tian F, Wang Q, Peiffer J, Li M et al (2012) GAPIT: genome association and prediction integrated tool. Bioinformatics 28:2397–2399

Lopes MS, Bovenhuis, H, van Son M, Nordbø Ø, Grindflek EH, Knol EF, Bastiaansen, JWM (2017) Using markers with large effect in genetic and genomic predictions, J Animal Sci 95(1):59–71

Low JW, Mwanga ROM, Andrade M, Carey E, Ball A (2017) Tackling vitamin A deficiency with biofortified sweetpotato in sub-Saharan Africa. Global Food Secur 14:23–30

Meuwissen TH, Hayes BJ and Goddard ME (2001) Prediction of total genetic value using genome-wide dense marker maps. Genetics 157:1819–1829

Michel S, Löschenberger F, Ametz C, Pachler B, Sparry E, and Bürstmayr H (2019) Simultaneous selection for grain yield and protein content in genomics-assisted wheat breeding. Theor Appl Genet 132:1745–1760

Mollinari M, Olokulu B, Pereira GDS, Khan A, Gemenet DC, Yencho C, Zeng Z-B (2020) Unraveling the hexaploid sweetpotato inheritance using ultra-dense multilocus mapping. G3: Genes, Genomes, Genetics 10(1):281–292

Morgante F, Huang W, Maltecca C, Mackay TFC (2018) Effect of genetic architecture on the prediction accuracy of quantitative traits in samples of unrelated individuals. Heredity 120:500–514

Mwanga ROM, Andrade MI, Carey EM, Low JW, Yencho GC, Grüneberg WJ (2017) Sweetpotato (Ipomoea batatas L.). In: Genetic Improvement of Tropical Crops. Campos H, Caligari PDS (Eds). Springer. p 181–218

Nakaya A, Isobe N. (2012). Will genomic selection be a practical method for plant breeding? Ann Bot 110(6):1303–1316

Nyine M, Uwimana B, Blavet N, Hřibová E, Vanrespaille H, Batte M, Akech V, Brown A, Lorenzen J, Swennen R, Doležel J (2018) Genomic prediction in a multiploid crop: Genotype by environment interaction and allele dosage effects on predictive ability in banana. Plant Genome 11:170090

Oliveira IDB, Resende Jr MFR, Ferrão LFV, Amadeu RR, Endelman JB, Matias Kirst M, Coelho ASG, Munoz PR (2019) Genomic prediction of autotetraploids; influence of relationship matrices, allele dosage and continuous genotype calls in phenotype prediction. G3: Genes Genomes Genetics 9:1189–1198

Pereira GDS, Gemenet DC, Mollinari M, Olukolu BA, Diaz F, Mosquera V, Gruneberg WJ, Khan A, Yencho GC, Zeng Z-B (2019) Multiple QTL mapping in autopolyploids: a random-effect model approach with application in a hexaploid sweetpotato full-sib population. BioRxiv Preprint doi: https://doi.org/10.1101/622951

Poland JA, Rife TW (2012) Genotyping-by-sequencing for plant breeding and genetics. Plant Genome J 5:92–102

Poplin R, Ruano-Rubio V, DePristo MA, Fennell TJ, Carneiro MO, Van der Auwera GA, Kling DE, et al (2017) Scaling accurate genetic variant discovery to tens of thousands of samples. BioRxiv Preprint. DOI:10.1101/201178

Raman H, Raman R, Kilian A, Detering F, Carling J, et al (2014) Genome-wide delineation of natural variation for pod shatter resistance in Brassica napus. PLoS ONE 9(7): e101673

Slater AT, et al (2016) Improving genetic gain with genomic selection in autotetraploid potato. The Plant Genome 9(3):1–15

Spindel JE, Begum H, Akdemir D, Collard B, Redoña E, Jannink J-L, McCouch S (2016) Genome-wide prediction models that incorporate de novo GWAS are a powerful new tool for tropical rice improvement. Heredity 116:395–408

Uitdewilligen JG, AM Wolters BB, D’hoop TJ, Borm RG, Visser et al (2013) A next-generation sequencing method for genotyping-by-sequencing of highly heterozygous autotetraploid potato. PLoS One 8: e62355

VanRaden PM (2008) Efficient methods to compute genomic predictions. Journal of Dairy Sci 91(11):4414–4423

Varona L, Legarra A, Toro MA, Vitezica ZG (2018) Non-additive Effects in Genomic Selection. Front Genet 9:78

Velazco JG, Malosetti M, Hunt CH, Mace ES, Jordan DR, van Eeuwijk FA (2019) Combining pedigree and genomic information to improve prediction quality: an example in sorghum. Theor Appl Genet 132:2055–2067

Vitezica ZG, et al (2013) On the additive and dominant variance and covariance of individuals within the genomic selection scope. Genetics 195:1223–1230

Wadl PA, Olukolu BA, Branham SE, Jarret RL, Yencho GC, et al (2018) Genetic Diversity and Population Structure of the USDA Sweetpotato (Ipomoea batatas) Germplasm Collections Using GBSpoly. Front Plant Sci 9:1–13

Wang X, Xu Y, Hu Z, Xu C (2018) Genomic selection methods for crop improvement: Current status and prospects. The Crop Journal 6:330–340

Watson A, Ghosh S, Williams M. et al (2018) Speed breeding is a powerful tool to accelerate crop research and breeding. Nature Plants 4:23–29 (2018)

Wu S, Lau KH, Cao Q, Hamilton JP, Sun H, Zhou C et al (2018) Genome sequences of two diploid wild relatives of cultivated sweetpotato reveal targets for genetic improvement. Nature communications 9:4580

Xu Y, Crouch JH (2008) Marker-assisted selection in plant breeding: from publications to practice. Crop Sci 48:391–407

Zhang Z, Ersoz E, Lai CQ, Todhunter RJ, Tiwari HK et al (2010) Mixed linear model approach adapted for genome-wide association studies. Nat Genet 42:355–360

