## Supplementary figures and images for "Sequencing depth and genotype quality: Accuracy and breeding operation considerations for genomic selection applications in autopolyploid crops"

### Online Resource 1

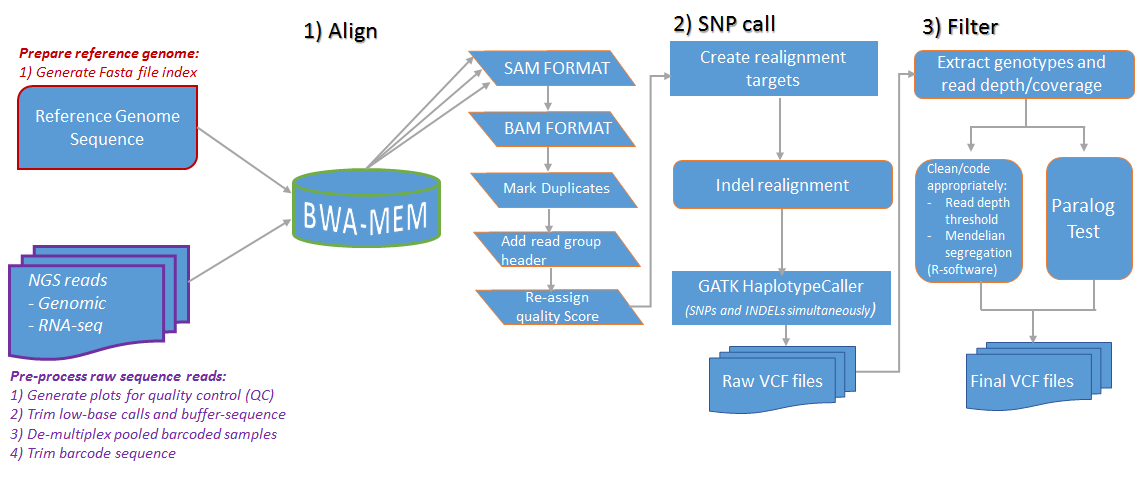
